# Immunogenicity of convalescent and vaccinated sera against clinical isolates of ancestral SARS-CoV-2, beta, delta, and omicron variants

**DOI:** 10.1101/2022.01.13.475409

**Authors:** Arinjay Banerjee, Jocelyne Lew, Andrea Kroeker, Kaushal Baid, Patryk Aftanas, Kuganya Nirmalarajah, Finlay Maguire, Robert Kozak, Ryan McDonald, Amanda Lang, Volker Gerdts, Sharon E. Straus, Lois Gilbert, Angel Xinliu Li, Mohammad Mozafarihasjin, Sharon Walmsley, Anne-Claude Gingras, Jeffrey L. Wrana, Tony Mazzulli, Karen Colwill, Allison J. McGeer, Samira Mubareka, Darryl Falzarano

## Abstract

The omicron variant of concern (VOC) of SARS-CoV-2 was first reported in November 2021 in Botswana and South Africa. Omicron has evolved multiple mutations within the spike protein and the receptor binding domain (RBD), raising concerns of increased antibody evasion. Here, we isolated infectious omicron from a clinical specimen obtained in Canada. The neutralizing activity of sera from 65 coronavirus disease (COVID-19) vaccine recipients and convalescent individuals against clinical isolates of ancestral SARS-CoV-2, beta, delta, and omicron VOCs was assessed. Convalescent sera from unvaccinated individuals infected by the ancestral virus during the first wave of COVID-19 in Canada (July, 2020) demonstrated reduced neutralization against beta and omicron VOCs. Convalescent sera from unvaccinated individuals infected by the delta variant (May-June, 2021) neutralized omicron to significantly lower levels compared to the delta variant. Sera from individuals that received three doses of the Pfizer or Moderna vaccines demonstrated reduced neutralization of the omicron variant relative to ancestral SARS-CoV-2. Sera from individuals that were naturally infected with ancestral SARS-CoV-2 and subsequently received two doses of the Pfizer vaccine induced significantly higher neutralizing antibody levels against ancestral virus and all VOCs. Importantly, infection alone, either with ancestral SARS-CoV-2 or the delta variant was not sufficient to induce high neutralizing antibody titers against omicron. This data will inform current booster vaccination strategies, and we highlight the need for additional studies to identify longevity of immunity against SARS-CoV-2 and optimal neutralizing antibody levels that are necessary to prevent infection and/or severe COVID-19.

## INTRODUCTION

SARS-CoV-2 has continued to evolve since its emergence in December 2019 ^1,2^. Variants of SARS-CoV-2 that demonstrate potential for interference with diagnostics, therapies, and vaccine efficacy, along with evidence for increased transmissibility or disease severity are termed variants of concern (VOCs). The most recent VOC, omicron was first reported in November 2021 in Botswana and South Africa ^3,4^. The omicron variant has evolved multiple mutations within the spike protein and the receptor binding domain (RBD) that raise concerns regarding a possible increased ability to evade pre-existing antibodies, both from prior infection and from vaccination ^5^. The omicron variant has demonstrated increased transmission and a higher level of resistance to antibody-mediated neutralization ^4,5^. However, little is known about its pathogenicity and whether disease severity is altered in convalescent, vaccinated or unvaccinated individuals. In Canada, long-term care (LTC) residents were prioritized for third vaccine doses against SARS-CoV-2 based on the observation that antibody titers in older adults waned within six months of their second vaccine dose ^6,7^. The neutralizing potential of antibodies generated in LTC residents against VOCs, such as delta and omicron after three doses of mRNA vaccines remain unknown. Thus, to better assess the efficacy of antibody-mediated neutralization against ancestral SARS-CoV-2 and VOCs (beta, delta, and omicron) in naturally infected and vaccinated individuals, we collected sera from multiple cohorts and tested their neutralization ability against clinical isolates of ancestral SARS-CoV-2 and VOCs.

## RESULTS

### Isolation of viruses

Isolates of VOCs used in this study were derived from clinical specimens. Nasopharyngeal swabs were collected from PCR positive patients, and virus isolation was performed on African Green monkey kidney cells (Vero’76) as previously described ^8^. We confirmed the whole genome sequence of the isolates and determined their phylogenetic relationship with other SARS-CoV-2 isolates (Figure 1A and see supplementary Table S1). Beta, delta and omicron isolates used in this study aligned with their expected lineages (Figure 1A). SARS-CoV-2 can rapidly adapt in cell culture and evolve adaptive mutations. We confirmed mutations across the full-length viral genome, including the spike protein for all variants prior to using the viruses in a micro-neutralization assay (Figure 1B and see supplementary Table S2).

**Figure 1.**
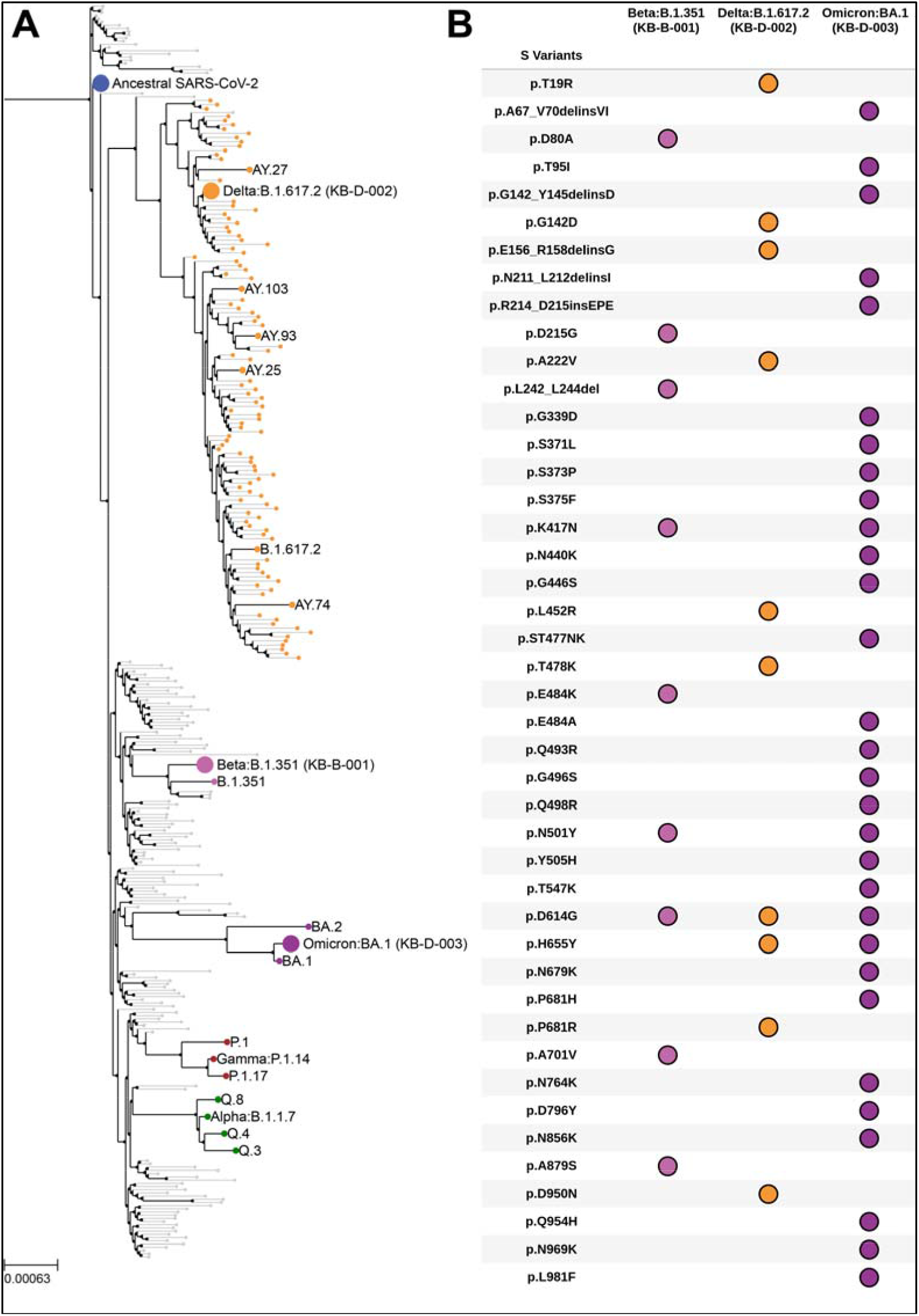
Spike mutations and phylogenetic analyses of clinical isolates of VOCs. **(A)** Clinical isolates of VOCs used in this study (Delta, B.1.617.2 KB-D-002; Beta, B.1.351 KB-B-001 and Omicron, BA.1 KB-D-003) were sequenced and assigned lineages by phylogeny analysis. **(B)** Mutations within the spike (S) protein of each variant are shown here. Delins, deletion+insertion; ins, insertion; p, amino acid position. See also supplementary Tables S1 and S2.

### Neutralization of omicron by convalescent sera from individuals infected with ancestral SARS-CoV-2 or the delta variant

To determine the neutralizing titer of sera from individuals that were naturally infected with SARS-CoV-2, we tested convalescent sera from individuals that were infected with ancestral SARS-CoV-2 during the first wave of coronavirus disease (COVID-19) in Canada (July 2020; see supplementary Table S3). Serum samples were collected 1-5 months after the onset of COVID-19 (see supplementary Table S3). Convalescent sera (n=15) from individuals infected with ancestral SARS-CoV-2 during the first wave of COVID-19 in Canada contained significantly lower neutralizing antibodies against both beta (p=0.0058) and omicron (p=0.0019) variants, relative to the ancestral virus (Figure 2A). However, neutralizing antibody titers in these serum samples were not significantly different between ancestral SARS-CoV-2 and the delta variant (p=0.1691; Figure 2A). Next, the neutralizing antibody titers in convalescent sera (n=10) from individuals who were infected with the delta variant in Canada between May and June, 2021 were determined (see supplementary Table S3). Serum samples were collected 1-2 months after the date of onset of COVID-19 (see supplementary Table S3). Convalescent delta sera contained lower levels of neutralizing antibodies against both beta (p=0.0468) and omicron variants (p=0.049) relative to delta variant (Figure 2B), while titers against ancestral SARS-CoV-2 and delta were comparable (p=0.6034; Figure 2B). These data suggest that infection with either ancestral SARS-CoV-2 or the delta variant induces cross-neutralizing antibody with comparable titers against both viruses. However, natural infection with either ancestral SARS-CoV-2 or the delta variant induces significantly lower levels of neutralizing antibodies against both beta and omicron variants.

**Figure 2.**
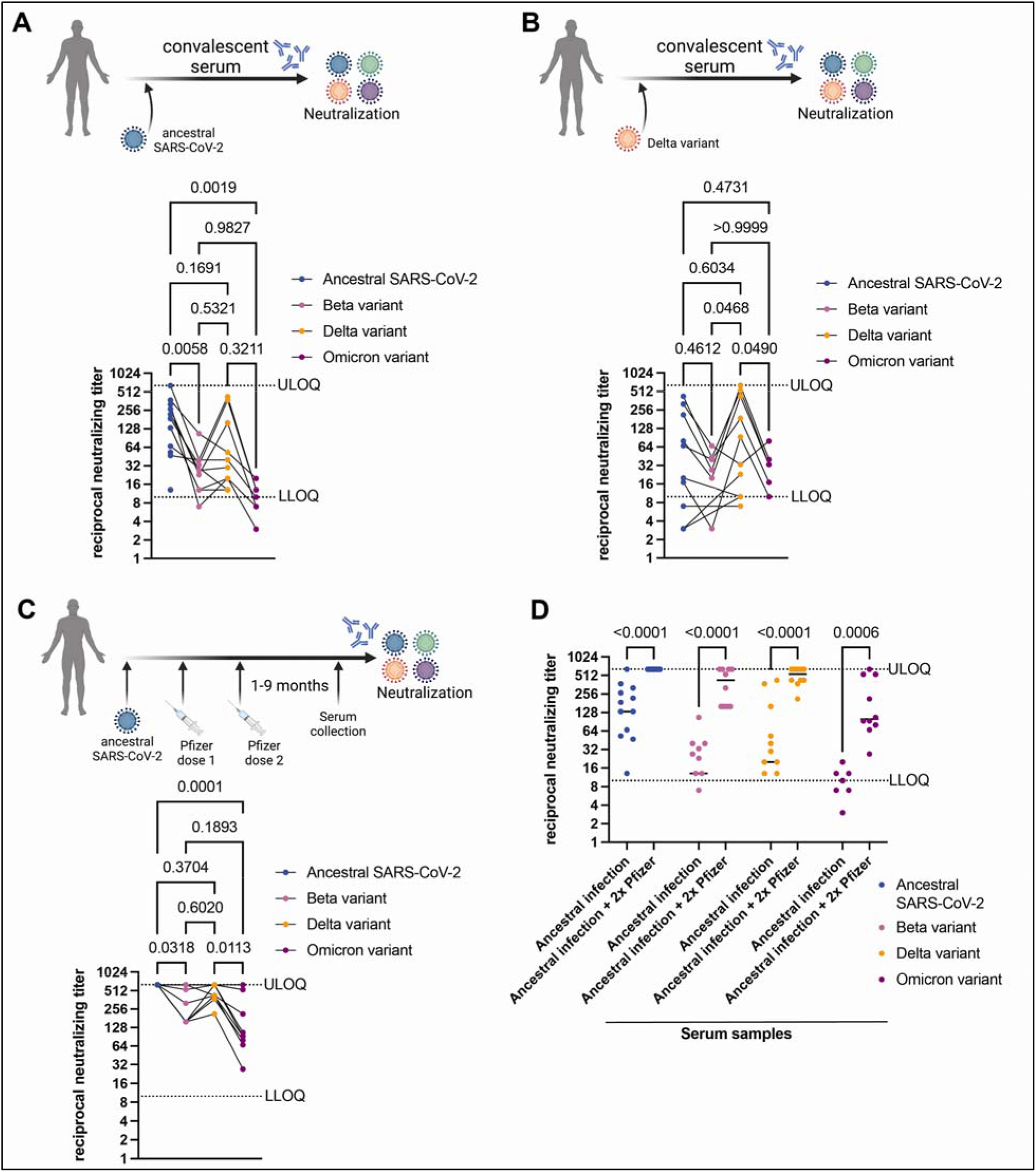
Detection of neutralizing antibodies in convalescent sera and sera from naturally infected individuals who subsequently received two doses of an mRNA vaccine. **(A)** Neutralizing antibody titers in convalescent sera collected from individuals that were infected with ancestral SARS-CoV-2 during the first wave in Canada (July 2020; n=15) tested against ancestral SARS-CoV-2, beta, delta and omicron variants. **(B)** Neutralizing antibody titers in convalescent sera collected from individuals that were infected with the delta variant of SARS-CoV-2 in Canada (May-June, 2021; n=10) tested against ancestral SARS-CoV-2, beta, delta and omicron variants. **(C)** Neutralizing antibody titers in sera collected from individuals that were infected with ancestral SARS-CoV-2, followed by two doses of the Pfizer BNT162b2 vaccine tested against ancestral SARS-CoV-2, beta, delta, and omicron variants (n=10). Neutralizing antibody titers against ancestral SARS-CoV-2 reached the upper limit of detection in our assay. **(D)** Neutralizing antibody titers in convalescent sera from individuals infected with ancestral SARS-CoV-2 compared to neutralizing antibody levels in sera from individuals who were infected with the ancestral virus and subsequently received two doses of the Pfizer mRNA vaccine. Data compiled and replotted from panels A and C for comparison. Mean values are indicated by horizontal black bars. Individual data points are shown and titers for matching serum samples are shown across different virus isolates. N = 15 or 10, *p* values are indicated in the figures (Tukey’s multiple comparisons test with alpha = 0.05 or Sidak’s multiple comparisons test with alpha = 0.05). Samples with neutralizing titer of 0 are not shown. LLOQ, lower limit of quantitation; ULOQ; upper limit of quantitation. See also supplementary Table S3.

### Neutralization of omicron by sera from individuals who received two doses of the Pfizer BNT162b2 mRNA vaccine post COVID-19 infection

Our data suggest that natural infection with ancestral SARS-CoV-2 is not sufficient to induce comparable levels of neutralizing antibodies against beta and omicron variants (Figure 2A). Next, we determined levels of neutralizing antibodies in sera (n=10) from individuals who had received two doses of the Pfizer BNT162b2 vaccine after being naturally infected with ancestral SARS-CoV-2 (see supplementary Table S3). Two doses of the Pfizer BNT162b2 vaccine after natural infection led to higher levels of neutralizing antibodies against ancestral SARS-CoV-2 that were significantly higher than levels against the beta (p=0.0318) and omicron (p=0.0001) variants (Figure 2C). Infection and subsequent two dose vaccination with Pfizer BNT162b2 induced higher levels of neutralizing antibodies against ancestral virus and all VOCs, including omicron than infected only individuals (Figure 2D).

### Neutralization of omicron by sera from individuals that received one dose of the Pfizer BNT162b2 mRNA vaccine

To determine neutralizing antibody titers in sera from individuals that received one dose of the Pfizer BNT162b2 mRNA vaccine, we collected sera (n=10) one month after the first dose of the vaccine and tested neutralizing antibody titers against ancestral SARS-CoV-2, beta, delta and omicron variants (see supplementary Table S3). Low neutralizing antibody titers against the ancestral virus were detected in some samples; however, no neutralization of the three VOCs was observed, with the exception of one serum sample that had detectable levels of neutralizing antibodies against all VOCs (Figure 3A).

**Figure 3.**
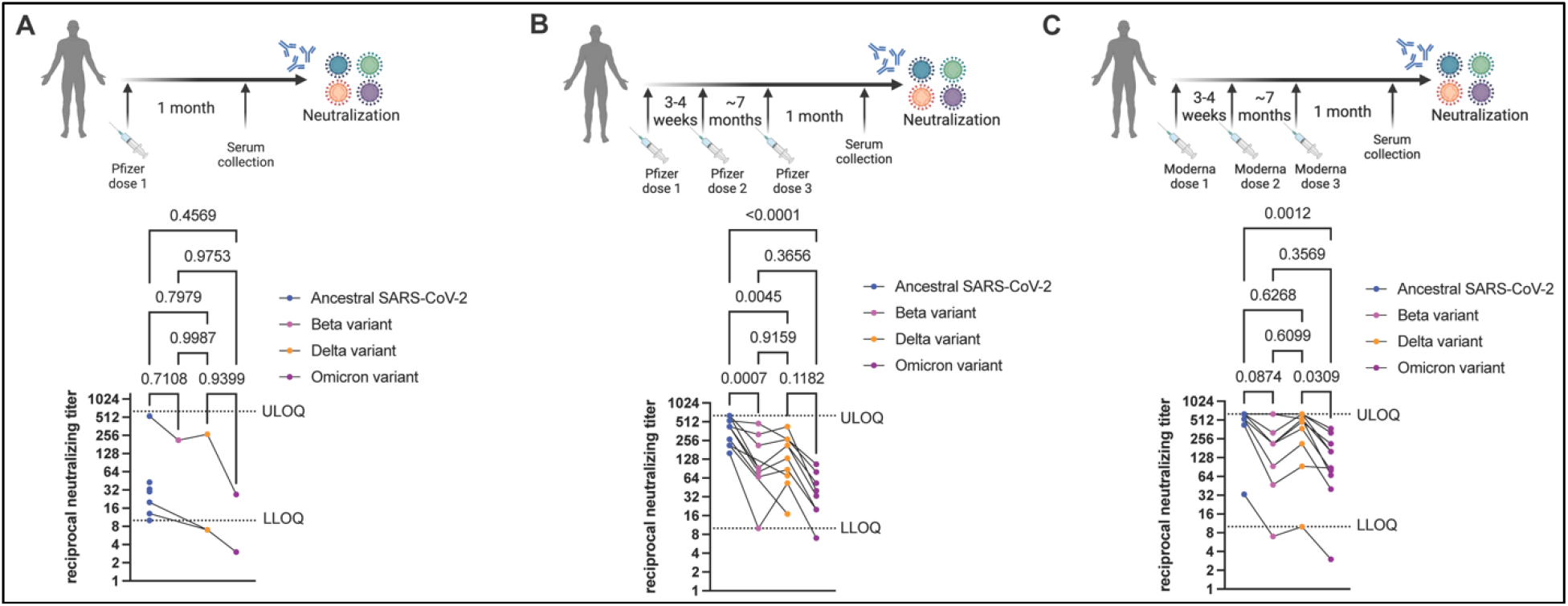
Detection of neutralizing antibodies in sera from vaccinated individuals. **(A)** Neutralizing antibody titers in sera collected from individuals that received one dose of the Pfizer BNT162b2 vaccine (n=10) tested against ancestral SARS-CoV-2, beta, delta, and omicron variants. **(B)** Neutralizing antibody titers in sera collected from individuals that received three doses of the Pfizer BNT162b2 vaccine (n=10) tested against ancestral SARS-CoV-2, beta, delta, and omicron variants. **(C)** Neutralizing antibody titers in sera collected from individuals that received three doses of the Moderna mRNA-1273 vaccine (n=10) tested against ancestral SARS-CoV-2, beta, delta and omicron variants. Individual data points are shown and titers for matching serum samples are shown across different virus isolates. N = 10, *p* values are indicated in the figures (Tukey’s multiple comparisons test with alpha = 0.05). Samples with neutralizing titer of 0 are not shown. LLOQ, lower limit of quantitation; ULOQ; upper limit of quantitation. See also supplementary Table S3.

### Neutralization of omicron by sera from triple vaccinated individuals

Additional booster vaccinations have been deemed critical in protecting us from VOCs in part by inducing higher levels of neutralizing antibodies. Thus, we tested the levels of neutralizing antibodies in sera collected from long-term care residents that received three doses of the Pfizer BNT162b2 (n=10) ^9^ or the Moderna mRNA-1273 (n=10) ^10^ vaccines (see supplementary Table S3). For both vaccine recipients, doses 1 and 2 were received 3-4 weeks apart. The third vaccine dose was received ~7 months after dose 2, and serum samples were collected 1 month after the third dose. Sera from individuals that received three doses of the Pfizer BNT162b2 vaccine induced high neutralizing titers against ancestral SARS-CoV-2, but levels of neutralizing antibodies were significantly lower against beta (p=0.0007), delta (p=0.0045) and omicron (p<0.0001) variants, compared to ancestral SARS-CoV-2 (Figure 3B). Sera from individuals that received three doses of the Moderna mRNA-1273 vaccine induced high neutralizing titers against ancestral SARS-CoV-2, but levels of neutralizing antibodies were significantly lower against the omicron variant, relative to ancestral SARS-CoV-2 (p=0.0012; Figure 3C). Neutralizing antibody titers were not significantly different against ancestral SARS-CoV-2, beta and delta variants. Serum samples from individuals that received 3x doses of the Pfizer BNT162b2 vaccine contained 2.86x, 2.25x and 10.3x lower mean neutralizing antibody titers against beta, delta and omicron VOCs, respectively, relative to ancestral SARS-CoV-2 (Figure 4). Serum samples from individuals that received 3x doses of the Moderna mRNA-1273 vaccine contained 1.7x, 1.26x and 3.48x lower mean neutralizing antibody titers against beta, delta and omicron VOCs, respectively, relative to ancestral SARS-CoV-2 (Figure 4).

**Figure 4.**
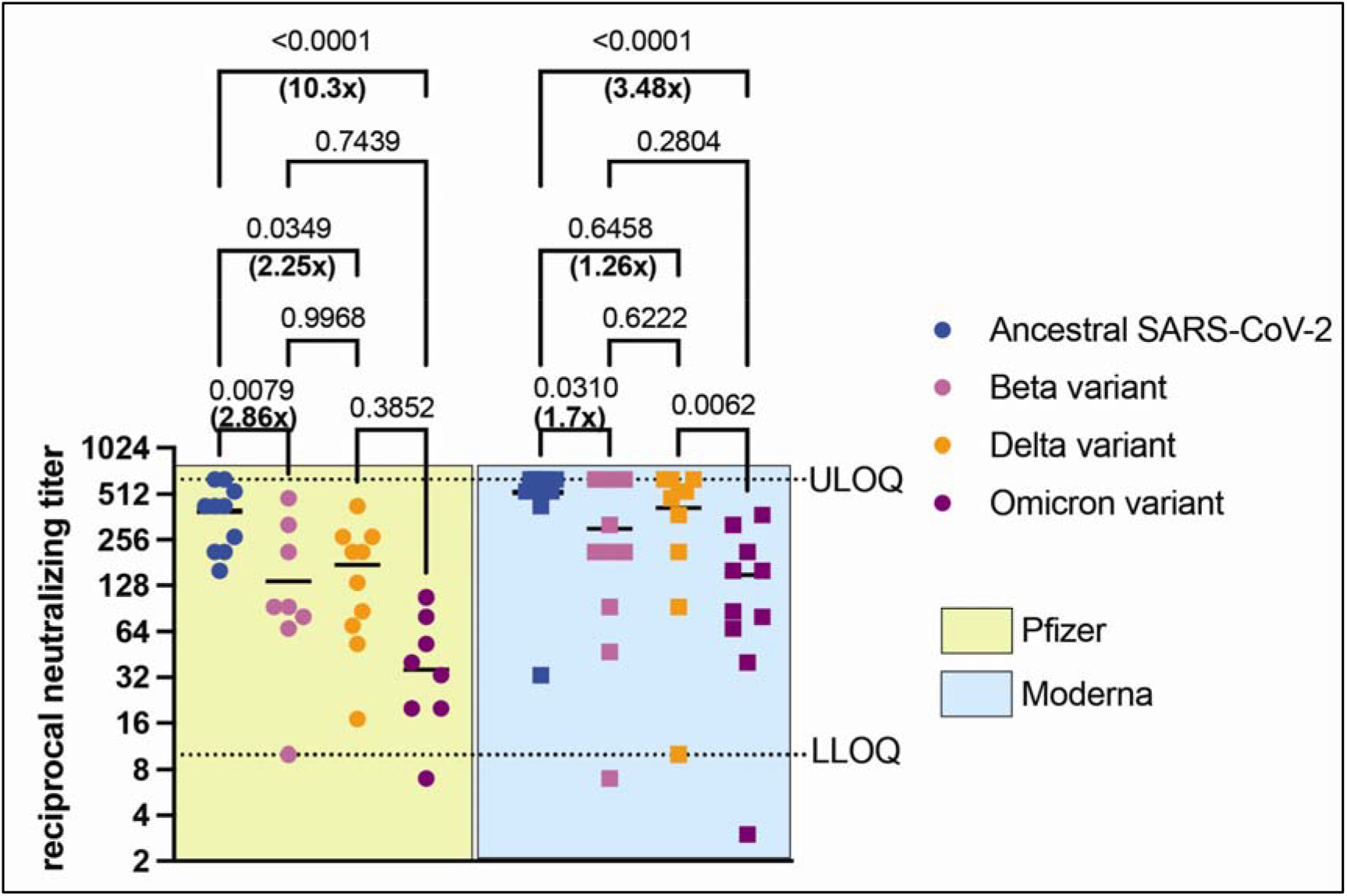
Neutralizing antibody levels in individuals vaccinated with 3x doses of the Pfizer BNT162b2 or Moderna mRNA-1273 vaccines against ancestral SARS-CoV-2 and VOCs. To compare neutralizing antibody titers in sera collected from long-term care residents that received three doses of either the Pfizer BNT162b2 or Moderna mRNA-1273 vaccines, we reanalyzed the data in Figure 3. Neutralizing antibody levels against ancestral SARS-CoV-2, beta, delta, and omicron variants are shown here. Mean values are indicated by horizontal black bars. Data are represented as mean, n = 15 or 10, *p* values are indicated in the figure (Sidak’s multiple comparisons test with alpha = 0.05). Fold reduction in mean neutralizing antibody titers against VOCs, relative to ancestral SARS-CoV-2 is indicated. Samples with neutralizing titer of 0 are not shown. LLOQ, lower limit of quantitation; ULOQ; upper limit of quantitation. See also supplementary Table S3.

## DISCUSSION

The emergence of an yet another SARS-CoV-2 VOC, omicron has led to increasing speculation about the ability of this variant to escape vaccine and natural infection-mediated immunity. The current generation of COVID-19 mRNA vaccines are designed using the *spike* gene sequence of ancestral SARS-CoV-2 ^9,10^. The omicron variant has accumulated 29 amino acid substitutions, 3 amino-acid deletions and a 3-residue insertion within the spike protein compared to the ancestral SARS-CoV-2 Wuhan isolate ^5^. Accumulating data suggest that the omicron variant is at least partially resistant to neutralization by antibodies in vaccinated individuals, along with partial or complete resistance to neutralization by therapeutic monoclonal antibodies ^5^. Emerging data demonstrate that T-cell-mediated immunity generated upon infection or vaccination likely remain effective against the omicron variant ^11^, and an additional booster vaccine dose results in higher levels of antibodies against the omicron variant when tested using pseudotyped viruses ^12^. Despite these recent advances, considerable gaps currently exist in our knowledge regarding the ability of omicron to cause severe COVID-19 and whether partial or complete escape of vaccine or natural infection-mediated immunity occurs and if escape is age dependent. In addition, it is not known if omicron has altered host range, or if transmissibility is increased and whether there are changes in cellular tropism. Furthermore, data on neutralizing antibody titers against clinical isolates of omicron are limited. Thus, as part of this study, we determined the levels of neutralizing antibodies in individuals that were naturally infected, infected and subsequently vaccinated, or vaccinated with three doses of mRNA vaccines using clinical isolates of ancestral SARS-CoV-2, beta, delta and omicron variants.

When omicron was first detected, multiple laboratories reported difficulties in isolating and generating laboratory stocks of this variant. In this study, we used Vero’76 cells to isolate the omicron variant from a clinical specimen (nasopharyngeal swab) that was collected from a Canadian patient. We also confirmed the whole genome sequences of the omicron variant, along with beta and delta variants (Figure 1 and Table 1). Thus, we report that Vero’76 cells are sufficient to facilitate the isolation and propagation of the omicron variant.

Next, we tested the levels of neutralizing antibody titers in convalescent sera against the ancestral virus, beta, delta and omicron variants (Figure 2). Infection with both the ancestral virus and the delta variant induced high levels of neutralizing antibodies against each other. However, the levels of neutralizing antibodies in convalescent sera against the omicron variant were lower compared to both the ancestral virus and the delta variant (Figures 2A and 2B). Thus, our data suggest that infection alone, either with the ancestral virus or the delta variant may not be sufficient to induce high levels of neutralizing antibodies against the omicron variant. Indeed, our data demonstrate that two doses of the Pfizer BNT162b2 mRNA vaccine following infection with ancestral SARS-CoV-2 induced significantly higher levels of neutralizing antibodies against ancestral SARS-CoV-2 and beta, delta and omicron variants (Figure 2D).

Our data highlight that one dose of the Pfizer vaccine is not sufficient to induce high levels of neutralizing antibodies against ancestral virus or variants (Figure 3A), thus the second vaccine doses appears to be critically required to induce neutralizing antibodies. Three doses of either the Pfizer BNT162b2 or Moderna mRNA-1273 vaccine induced comparable neutralizing antibodies against the beta and omicron variants (Figures 3B and 3C). However, levels of neutralizing antibodies against omicron were significantly lower compared to ancestral SARS-CoV-2 in serum samples from individuals vaccinated with 3x doses of either of the mRNA vaccines (Figures 3B, 3C and 4).

In summary, our data demonstrate that infection alone, either with ancestral SARS-CoV-2 or the delta variant is not sufficient to induce high levels of neutralizing antibodies against omicron. However, two doses of the Pfizer vaccine in previously infected individuals induces higher levels of neutralizing antibodies. While we did not test the effect of two doses of the Moderna mRNA-1273 vaccine in previously infected and recovered individuals, we speculate that the results will be comparable to the Pfizer BNT162b2 vaccine. Our data also show that while 3x doses of both mRNA vaccines induce neutralizing titers against omicron variant in long-term care residents, the levels of neutralizing antibodies remain significantly lower compared to ancestral SARS-CoV-2. Thus, our data support the ongoing third vaccine dose booster strategy for long-term care residents in Canada. Indeed, there is a need for studies to assess the optimal levels of neutralizing antibodies that are required for protection against infection and/or severe COVID-19, which will inform policies around vaccine boosters and enable equitable distribution of vaccines to end the ongoing pandemic.

## ACKNOWLEDGEMENTS

SARS-CoV-2 research is supported in the laboratory of D.F. by the Canadian Institutes of Health Research (CIHR; OV5-170349, VRI-173022 and VS1-175531). A.B. receives funding from VIDO. VIDO receives operational funding from the Government of Saskatchewan through Innovation Saskatchewan and the Ministry of Agriculture and from the Canada Foundation for Innovation through the Major Science Initiatives for its CL3 facility. Studies from which clinical samples were collected were funded by CIHR grants to A.J.M and S.M. (#439999, #465038); from the Canadian COVID-19 Immunity Task Force to S.E.S and A.J.M.; and from the Toronto COVID Action Initiative Fund from the University of Toronto to J.L.W. and T.M. A.B., A-.C.G., J.L.W., S.M. and D.F. are members of the CIHR-funded Coronavirus Variants Rapid Response Network (CoVaRR-Net). R.K. is supported by an Ontario Together grant. We acknowledge contributions by Dr. Andrew G. McArthur who connected our teams to facilitate virus sequencing. We acknowledge the help of Dr. Akarin Asavajaru who handled shipping and receiving of samples.

## AUTHOR CONTRIBUTIONS

Conceptualization, A.B., F.M., S.M. and D.F.; Sample Collection and Selection: S.E.S, L.G., A.X.L., M.M., S.W. and A.C.G; Methodology, A.B., J.L., A.K., K.B., P.A., F.M. and D.F.; Formal analysis, A.B., J.L., A.K. and F.M.; Reagents, J.L., R.K., R.M., A.L., J.L.W., T.M., A.J.M and S.M.; Funding acquisition, A.B. and D.F.; Writing – reviewing and editing, A.B., J.L., A.K., F.M., V.G., S.M. and D.F.; All authors reviewed the final manuscript; Supervision, A.B. and D.F.

## SUPPLEMENTARY TABLES

**Table S1.** Metadata for phylogenetic analysis

**Table S2.** Frequency of mutations (percentage read support) across the full genome of clinical isolates of beta, delta and omicron variants used in this study

**Table S3.** Serum samples from different cohorts

## METHODS

### RESOURCE AVAILABILITY

#### Lead Contact

Further information and requests for resources and reagents should be directed to and will be fulfilled by lead contacts, Drs. Darryl Falzarano and Arinjay Banerjee.

#### Materials availability

This study generated multiple virus isolates. The reagents will be made available on request through institutional Material Transfer Agreements for organizations that have a compliant BSL3 laboratory.

#### Data and code availability

All data and scripts associated with analyses of the virus sequences can be found here: https://github.com/fmaguire/voc_neutralisation_sc2_phylogenomics and DOI 10.5281/zenodo.5817727. Full sequences of the viral isolates have been deposited to NCBI BioProject PRJNA794206.

### EXPERIMENTAL MODEL

#### Cells and viruses

Vero’76 cells (CRL-1587, ATCC) were used to isolate and/or propagate all virus isolates using a previously published protocol ^8^. Briefly, the fluid from PCR-positive nasopharyngeal swabs received from Sunnybrook Research Institute (R.K, S.M. – ancestral SARS-CoV-2), omicron was identified by SPAR-Seq (PMID: 33658502) at the joint MSH/UHN Microbiology clinical diagnostic laboratory (J.L.W., T.M., S.M. – omicron) and beta at the Roy Romanow Provincial Laboratory (A.L., R.M.) were centrifuged at 8000x*g* for 15 minutes and 50μl removed and mixed with vDMEM containing 1μg/ml of TPCK trypsin. The mixture was added to a 24 well plate of Vero’76 cells and centrifuged for 1 h at 37°C at 800xg and then placed at 37°C for 30 min. The inoculum was removed and replaced with fresh vDMEM containing 1μg/ml of TPCK trypsin. Cell were monitored daily for cytopathic effect and on day 3 or 4, supernatant was passaged to fresh Vero’76 cells in a 6 well plate. Supernatant was subsequently collected on day 3 or 4 and passaged to T175 flasks to generate a p.1 virus stock. Virus stocks were subsequently titered on Vero’76 cells by TCID_50_ assay. Delta was obtained as a virus stock from the National Microbiology Laboratory and used to generate a stock as described.

For ancestral SARS-CoV-2, we used SARS-CoV-2/VIDO-1, the sequence for which has been previously reported (>hCoV-19/Canada/ON_ON-VIDO-01-2/2020|EPI_ISL_425177|2020-01-23). All work with infectious SARS-CoV-2 isolates were performed in a containment level 3 laboratory at the Vaccine and Infectious Disease Organization using approved protocols. Use of clinical specimen for virus isolation and use of human serum samples for micro-neutralization assays were approved by the University of Saskatchewan’s Biomedical Research Ethics Board (REB# 2591).

#### Serum samples

Serum samples were acquired from a series of different cohorts (see supplementary Table S3). Cohort participants provided informed consent for sharing of serum, and studies were approved by the Sunnybrook Research Institute (REB# 149-1994) and/or the Mount Sinai Hospital (REB# 02-0118-U, 20-0339-E, and 21-0069-E) Research Ethics Board ^13^. For samples from each cohort, samples were selected to have a representative range of anti-spike trimer and anti-RBD antibodies as measured by enzyme-linked immunosorbent assay ^14^.

### METHOD DETAILS

#### Sequencing and bioinformatic analyses

cDNA was synthesized from extracted RNA. In brief, 4 μL LunaScript RT SuperMix 5X (New England Biolabs, NEB, USA) and 8 μL nuclease free water, were added to 8 μL extracted RNA. cDNA synthesis was performed using the following conditions: 25 °C for 2 min, 55 °C for 20 min, 95 °C for 1 min, and holding at 4 °C.

Amplicons were generated from cDNA using ARTIC V4 primer pools (https://github.com/artic-network/artic-ncov2019). Two multiplex PCR tiling reactions were prepared by combining 2.5 μL cDNA with 12.5 μL Q5 High-Fidelity 2X Master Mix (NEB, USA), 6μL nuclease free water, and 4 μL of respective 10 μM ARTIC v4 primer pool (Integrated DNA Technologies). PCR cycling was then performed in the following manner: 98 °C for 30 s followed by 35 cycles of 98 °C for 15 s and 63 °C for 5 min.

Both PCR reactions were combined and cleaned with 1X ratio Sample Purification Beads (Illumina) at a 1:1 bead to sample ratio. The quantity of amplicons was measured with the Qubit 4.0 fluorometer using the 1X dsDNA HS Assay Kit (Thermo Fisher Scientific, USA) and the sequencing libraries were prepared using the Nextera DNA Flex Prep kit (Illumina, USA) as per manufacturer’s instructions. Paired-end (2×150 bp) sequencing was performed on a MiniSeq with a 300–cycle reagent kit (Illumina, USA) with a negative control library with no input SARS-CoV-2 RNA extract.

Raw reads underwent adapter/quality trimming (trim-galore v0.6.5) ^15^, host filtering and read mapping to reference [bwa v0.7.17 ^16^, samtools v.1.7 ^17,18^] trimming of primers (iVar v1.3 ^19^) and variant/consensus calling (freebayes v1.3.2 ^20^) using the SIGNAL workflow (https://github.com/jaleezyy/covid-19-signal) v1.4.4dev (#60dd466) with the ARTICv4 amplicon scheme (from https://github.com/artic-network/artic-ncov2019) and the MN908947.3 SARS-CoV-2 reference genome and annotations. Additional quality control and variant effect annotation (SnpEff v5.0-0 ^21^) was performed using the ncov-tools v1.8.0 (https://github.com/jts/ncov-tools/). Finally, PANGO lineages were assigned to consensus sequences using pangolin v3.1.17 (with the PangoLEARN v2021-12-06 models) ^22^, scorpio v0.3.16 (with constellations v0.1.1) [citation: https://github.com/cov-lineages/scorpio], and PANGO-designations v1.2.117 ^23^. Variants were summarised using PyVCF v0.6.8 [citation: https://github.com/jamescasbon/PyVCF] and pandas v1.2.4 ^24^. Phylogenetic analysis was performed using augur v13.1.0 ^25^ with IQTree (v2.2.0_beta) ^26^ and the resulting phylogenetic figure generated using ETE v3.1.2 ^27^. Contexual sequences were incorporated into the phylogenetic analysis by using Nexstrain’s ingested GISAID metadata and pandas to randomly sample a representative subset of sequences (jointly deposited in NCBI and GISAID) that belonged to lineages observed in Canada (see also Supplemental Tables S1 and S2 for metadata).

#### Micro-neutralization assay

Serum samples were heat-inactivated at 56°C for 30 minutes and then serially diluted 1:2 in DMEM supplemented with 2% FBS and 1% P/S (vDMEM) in 96-well blocks. The final volume of diluted serum per well was 180 μL. Each virus was diluted to 500 TCID_50_/mL (25 TCID_50_ per well) in vDMEM, from which 180 μL was added to each well containing serum. Positive controls containing only media and negative controls containing only virus were also included. The serum-virus mixture was incubated at 37°C for 1 h and then added to cultured Vero’76 cells in 96 well plates in triplicate. After five days the cells were evaluated for cytopathic effect (CPE), and the endpoint neutralization titer (the highest dilution of sera without CPE) was recorded. The final neutralization titer is reported as the average of the three replicates per sample. As control, each virus isolate that was diluted for our micro-neutralization assay was also back-titered using 2-fold serial dilutions.

#### Statistical analysis

Statistical analyses were performed using Tukey’s multiple comparisons test with alpha = 0.05 or Sidak’s multiple comparisons test with alpha = 0.05.

